# Single-cell analysis of bovine muscle-derived cell types for cultured meat production

**DOI:** 10.1101/2022.09.02.506369

**Authors:** Tobias Messmer, Richard GJ Dohmen, Lieke Schaeken, Lea Melzener, Rui Hueber, Mary Godec, Mark J Post, Joshua E Flack

## Abstract

‘Cultured’ meat technologies leverage the proliferation and differentiation of animal-derived stem cells *ex vivo* to produce edible tissues for human consumption in a sustainable fashion. However, skeletal muscle is a dynamic and highly complex tissue, involving the interplay of numerous mono- and multinucleated cells, including muscle fibres, satellite cells (SCs) and fibro-adipogenic progenitors (FAPs), and recreation of the tissue *in vitro* thus requires the characterisation and manipulation of a broad range of cell types. Here, we use a single-cell RNA sequencing approach to characterise cellular heterogeneity within bovine muscle and muscle-derived cell cultures over time. Using this data, we identify numerous distinct cell type, and develop robust protocols for the easy purification and proliferation of several of these populations. We note overgrowth of undesirable cell types within heterogeneous proliferative cultures as a barrier to efficient cultured meat production, and use transcriptomics to identify conditions that favour the growth of SCs in the context of serum-free medium. Combining RNA velocities computed in silico with time-resolved flow cytometric analysis, we characterise dynamic subpopulations and transitions between active, quiescent, and committed states of SCs, and demonstrate methods for modulation of these states during long-term proliferative cultures. This work provides an important reference for advancing our knowledge of bovine skeletal muscle biology, and its application in the development of cultured meat technologies.

## Introduction

‘Cultured’ or ‘cultivated’ meat is an emergent technology that leverages in vitro proliferation and differentiation of stem cells to produce edible tissues that mimic conventional meat^1,2^. Whilst there are numerous potential advantages to this technology, including reduced greenhouse gas emissions and improved animal welfare^3^, many technical challenges remain, such as the removal of animal-derived components, scaling of culture volumes, and cost reduction^4,5^. Moreover, consumer acceptance will be reliant on the taste and texture of cultured meat products closely replicating those of the conventionally-reared equivalent^6,7^.

Meat is composed primarily of skeletal muscle, a complex tissue whose function requires the interplay of numerous mono- and multinucleated cell types, including muscle fibres, satellite cells (SCs) and fibro-adipogenic progenitors (FAPs), as well as vascular, nervous and connective tissue^8,9^. Whilst most cultured meat products currently being pursued consist of unstructured muscle fibres, with or without fat tissue, accurate recreation of the entire tissue requires the identification, purification, proliferation and characterisation of a broad range of cell types^10,11^. Descriptions of the composition of human and murine skeletal muscle at the cellular level are now available^12–16^, but a similarly detailed understanding of muscle biology in agriculturally relevant species, such as cattle, is currently lacking. Furthermore, the extent to which complex muscle-derived cell behaviours and interactions are recapitulated during in vitro culture is unclear^17,18^, limiting the progress of cultured meat development.

Here, we used droplet-based single-cell RNA sequencing (scRNA-seq) to profile bovine skeletal muscle, and muscle-derived cell cultures, in a time-resolved fashion across the process of cultured meat production. We use the resultant dataset to gain insight into transcriptional heterogeneity between and within cell types, and to inform various steps of cultured meat bioprocess design.

## Results

### scRNA-seq identifies 11 distinct cell types in bovine muscle

In order to investigate heterogeneity between and within bovine muscle-derived cell types, we used the 10x Chromium platform for scRNA-seq to study gene expression at five timepoints in a primary adult stem cell-based cultured meat production process (Fig. 1a, Supplementary Table 1). A total of 36129 single-cell transcriptomes, with an average expression of 3815 genes and 21131 transcripts per cell, were analysed across 10 donor cattle and assembled into a single dataset (Supplementary Fig. 1). Using UMAP dimensionality reduction, based on the first 30 principal components (Fig. 1b), cells from Timepoints 1 (muscle tissue) and 2 (passage 0, after 72 h in vitro culture) clustered separately from each other, and from Timepoints 3 to 5 (after passages 2, 5 and 8 respectively), indicating significant transcriptomic changes between timepoints and within cell types during the proliferative process.

**Figure 1:**
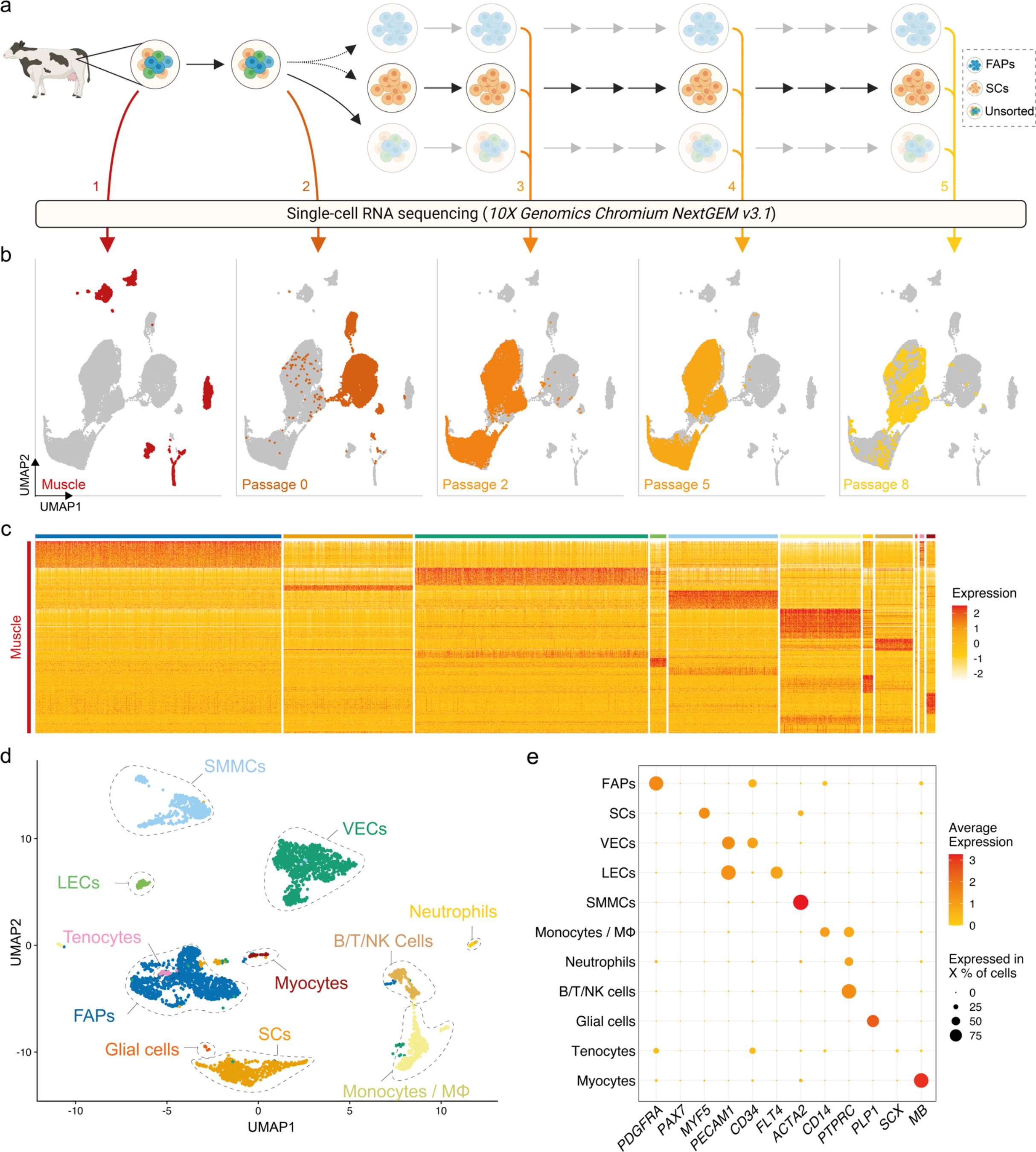
Single-cell RNA sequencing of bovine muscle and muscle-derived cell cultures. a) Overview of experimental design; coloured arrows and numerals indicate timepoints of RNA sequencing of single cells isolated from bovine muscle (Timepoint 1), at passage 0 after 72 h in serum-free growth medium (2) and at passages 2, 5 and 8 (3, 4 and 5 respectively); solid arrows indicate passaging, dotted arrows indicate FACS sorting; b) Combined UMAP plots showing single cells (individual points) from all five experimental timepoints; clustering is based on first 30 principal components, cells from each timepoint are coloured in each respective plot; c) Heatmap displaying normalised expression of significantly upregulated genes in each cluster identified in Timepoint 1. d) UMAP of Timepoint 1; cells are coloured and clusters labelled for their assigned cell types; e) Dotplot showing normalised expression of cell-type specific genes, averaged within each population; dot size indicates percentage of cells expressing the respective gene with at least 1 count.

Within enzymatically digested bovine muscle tissue (Timepoint 1), we identified 11 defined populations of mononuclear cells with distinct gene expression profiles (Fig. 1c; Supplementary Table 2), which were present in varying proportions in all donor animals (Supplementary Fig. 2a). Based on differential gene expression, these clusters were assigned to cell types previously characterised in other species^12,13,19^, namely fibro-adipogenic progenitors (FAPs), satellite cells (SCs), vascular and lymphatic endothelial cells (ECs; vascular, VECs; lymphatic, LECs, Supplementary Fig. 2b), smooth muscle and mesenchymal cells (SMMCs), monocytes/macrophages (Mϕ), neutrophils, lymphocytes (B/T/NK cells), glial cells, tenocytes and myocytes (Fig. 1d, Supplemental Fig. 2d). The expression of canonical genes in each of these populations in analogous studies indicated their conservation between mouse and cattle (Fig. 1e, Supplemental Fig. 2e)^14^.

### Cell types can be identified in vitro by unique surface marker expression

In order to identify proliferative muscle tissue-derived cell types that might be used for cultured meat production, we analysed the adherent cell fraction after 72 h (Timepoint 2) culture in serum-free growth medium (SFGM, Supplementary Table 3). We discerned 7 populations present in these cultures, which were separated from the same populations in Timepoint 1 by UMAP, indicating stark transcriptional changes during the transition to in vitro culture (Fig. 2a). GO terms corresponding to differentially expressed genes (independent of cell type) between Timepoints 1 and 2 (Fig. 2b) suggest that these transcriptional switches relate to decreased interaction with the extracellular matrix (ECM) and increased protein production, concomitant with increased cellular proliferation in the in vitro environment.

**Figure 2:**
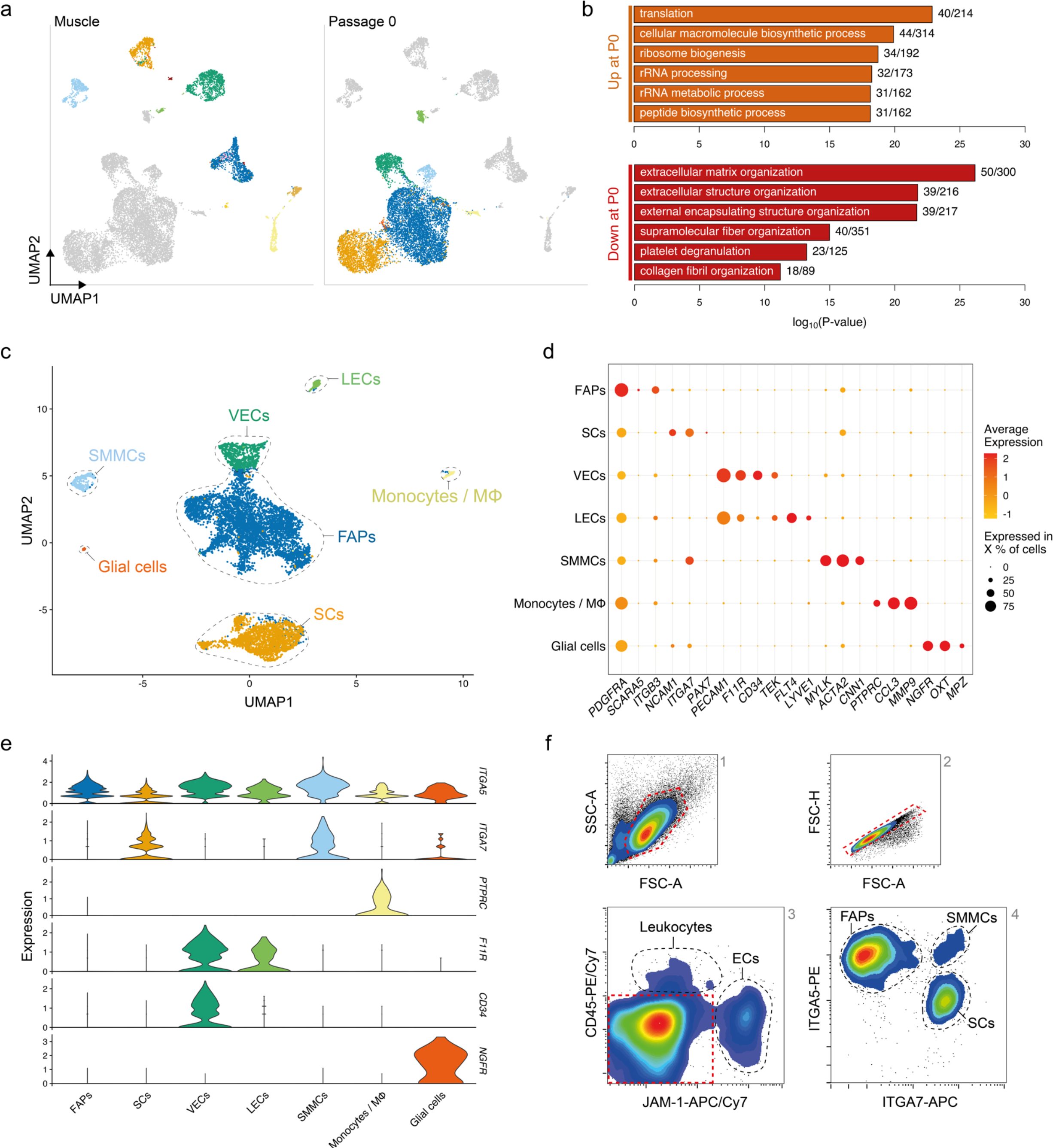
Transcriptomic analysis defines adherent cell types with distinct immunosurface phenotypes. a) Combined UMAP of Timepoints 1 and 2, coloured for populations in Timepoint 1 (left) or 2 (right), respectively; b) Top five most significantly enriched GO terms corresponding to differentially expressed genes up- (top) or downregulated (bottom) between Timepoints 1 and 2, after regressing out the effect of cell type; c) UMAP of Timepoint 2; cells are coloured and clusters labelled for their assigned cell types; d) Normalised gene expression of canonical markers in each population at Timepoint 2; e) Violin plots showing expression of surface markers in different cell types at Timepoint 2; f) Representative flow cytometry contour plots showing gating strategy for purification of cell types; red dashed lines indicate gates for the subsequent plot (from 1 to 4); black dashed lines indicate sorting gates for the labelled populations.

Comparison of Timepoints 1 and 2 enabled the assigning of cell identities to individual clusters (Fig. 2c), indicating that of the 11 populations identified in Timepoint 1, only myocytes, tenocytes, neutrophils and lymphocytes did not persist in short-term serum-free culture. Conversely, FAPs, SCs, SMMCs, ECs, glial cells and monocytes/macrophages remained present in cultures from all genotypes (donor animals) sequenced (Supplementary Figs. 3a, b). These same cell types were identified when culturing cells in growth medium (GM) containing 20% foetal bovine serum (FBS; Supplementary Fig. 3c, Supplementary Table 3), indicating that our SFGM formulation was able to support culture of muscle-derived cells as robustly as traditional serum-containing media.

Analysis of differentially expressed genes yielded highly and exclusively expressed markers for each population (Fig. 2d)^10,20^ that have previously been described in mouse and human^12–14,21^. Filtering these lists for genes encoding plasma membrane-localised proteins facilitated the identification of candidate cell surface markers for separation of populations by flow cytometry (Fig. 2e). Staining of muscle tissue-derived cells 72 h post-isolation for JAM-1 (*F11R*), CD45 (*PTPRC*), ITGA7 and ITGA5 confirmed the suitability of this quartet of markers as a FACS panel (Fig. 2f, Supplementary Fig. 3d).

### Four principal muscle-derived cell types can be proliferated in vitro

In order to characterise their in vitro phenotypes in more detail, we sorted muscle-derived cells into FAPs, SCs, ECs and SMMCs using the FACS panel previously described (Fig. 2). The purified populations exhibited markedly different morphologies; whilst FAPs had a spindle-like morphology and SCs were more spherical, ECs and SMMCs appeared flatter and larger (Fig. 3a). Flow cytometric analysis post-FACS using the same antibody panel confirmed high sorting purities for all populations (Fig. 3b). In addition, immunofluorescent staining for canonical markers PDGFRα, Pax7, TEK (also known as Tie2), and Calponin-1 (CNN1) further verified the identity of FAPs, SCs, ECs, and SMMCs respectively (Fig. 3c).

**Figure 3:**
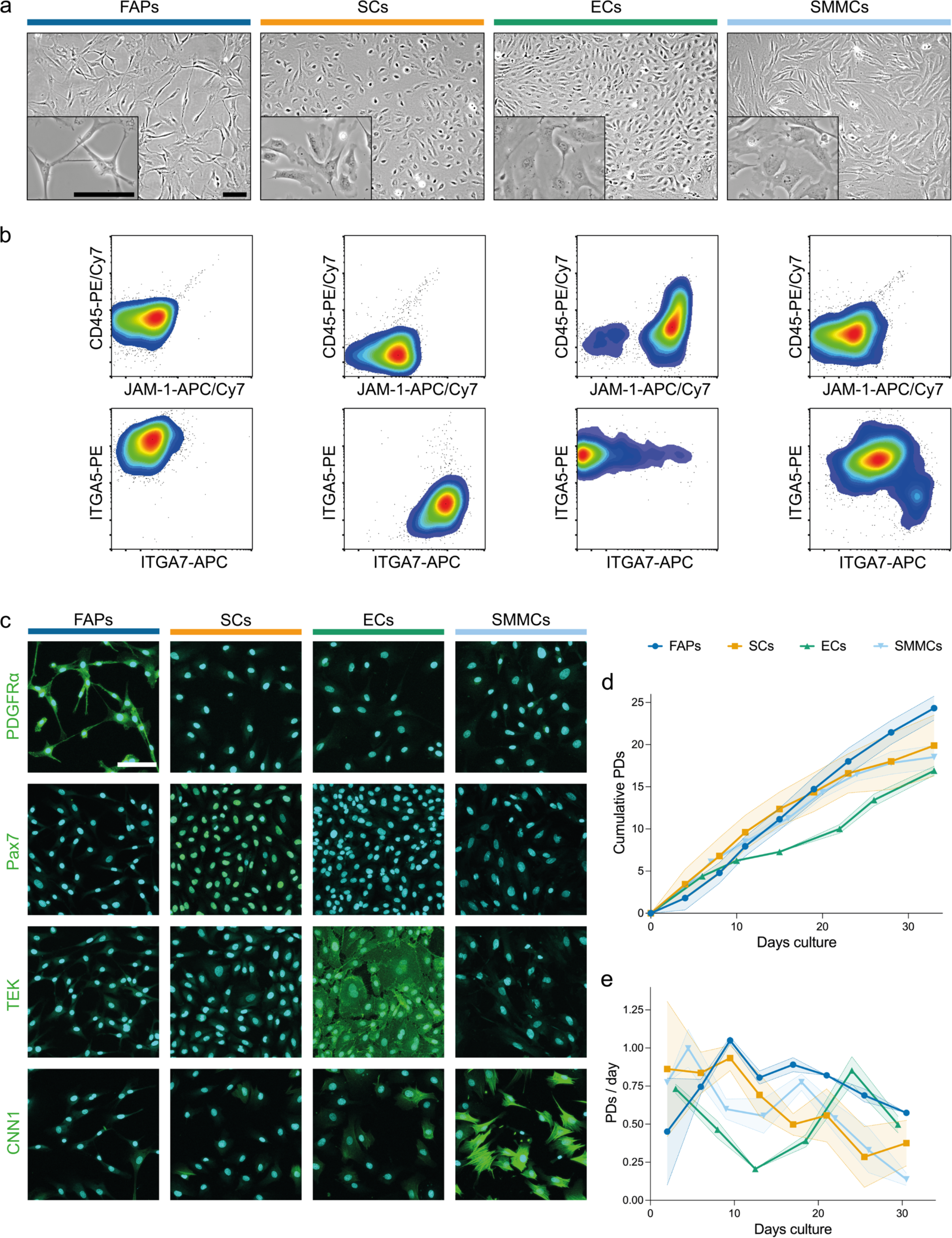
Four muscle-derived cell types can be purified and expanded in long-term culture. a) Brightfield microscopy images of purified FAPs (blue), SCs (orange), ECs (green) and SMMCs (light blue) in vitro; scale bars = 100 μm; b) Flow cytometry plots of purified cell types stained for CD45/JAM-1 (top) and ITGA5/ITGA7 (bottom); c) Immunofluorescent staining for PDGFRα, Pax7, TEK and CNN1 (green) in purified cell types; nuclei stained with Hoechst (cyan); scale bar = 100 μm; d) Long-term growth curves of purified cell types in vitro, shown as cumulative population doublings (PDs); shaded areas indicate standard deviation (*SD*), n = 3; e) Growth rates for long-term growth experiments shown in d), shown as PDs per day, n = 3.

Long-term proliferation of each of these populations was supported in SFGM (FAPs, SCs) or BioAMF-3 medium (ECs, SMMCs), with each culture undergoing at least 15 population doublings (PDs) over a period of at least 6 passages (Fig. 3d). Proliferation rates varied between cell types (for example, from 0.46 (ECs) to 1.0 PDs/day (SMMCs) at early passage), and tended to decrease over time (Fig. 3e).

### Optimised culture conditions prevent SC overgrowth by FAPs

We next aimed to understand heterogeneity within FACS-purified SC cultures during long-term proliferation. Surprisingly, we found that cells in these cultures clustered into two distinct populations, marked by differential expression of *ITGA7* (Fig. 4a). *ITGA7*- cells expressed FAP marker genes such as *ITGA5* (Supplemental Fig. 4a, Fig. 2), suggesting that these cells were FAPs that were inaccurately sorted into, and remained prevalent within, SC cultures. Quantification of these cell types showed that the proportion of FAPs increased over time, indicating that our standard culture conditions favoured long-term proliferation of FAPs over SCs (Fig. 4b, Supplemental Fig. 4b). The percentage of contaminating FAPs correlated negatively with myogenic differentiation, as determined by fusion index (Figs. 4c, d), emphasising the importance of eliminating overgrowth by non-SCs for cultured muscle production.

**Figure 4:**
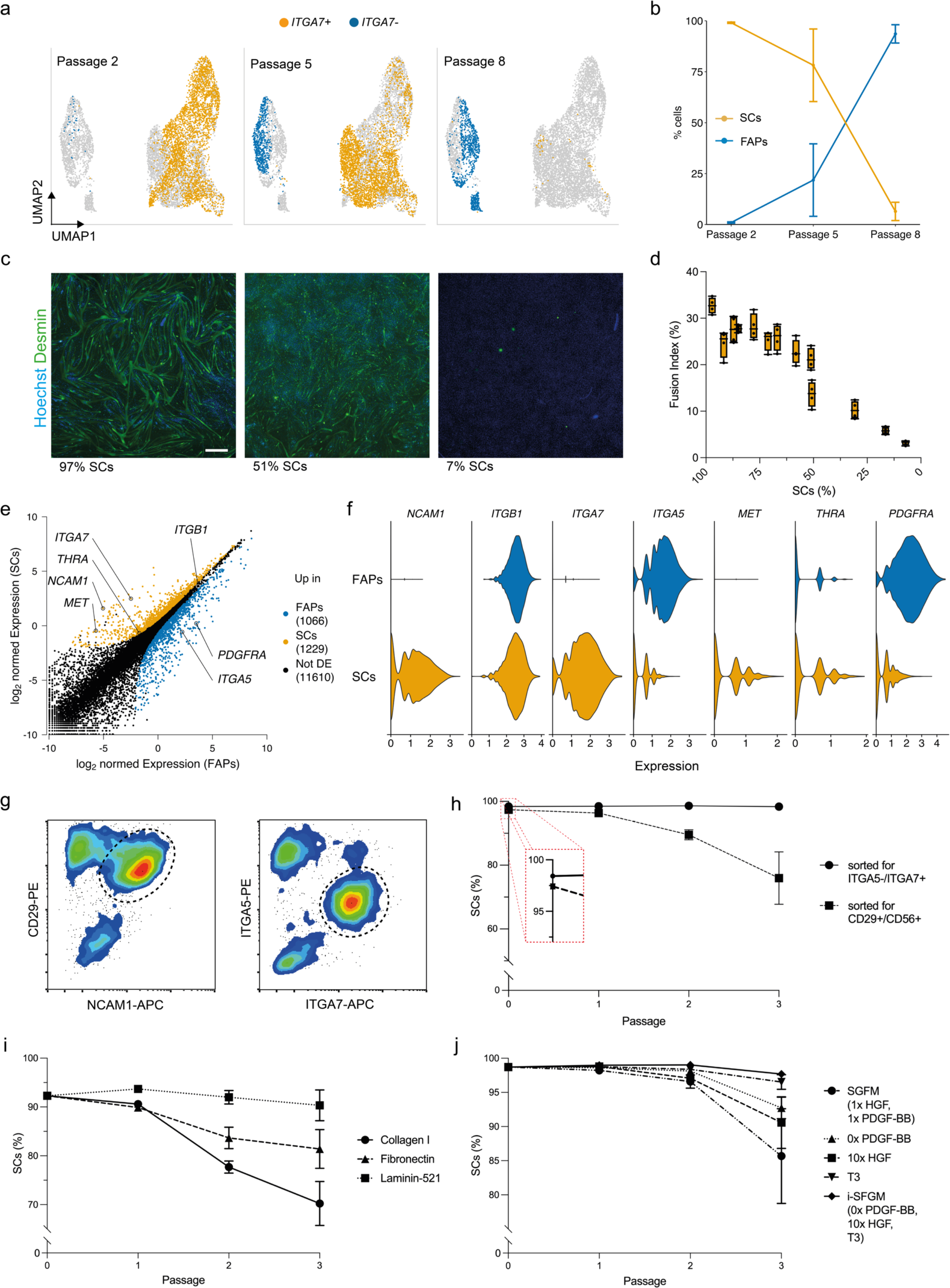
Optimised culture conditions prevent overgrowth of SCs. a) Combined UMAPs of sorted SCs at passages 2 (Timepoint 3, left), 5 (Timepoint 4, centre) and 8 (Timepoint 5, right); clusters in respective passages are coloured for *ITGA7* expression; b) Proportion of SCs (*ITGA7*+) during long-term culture; error bars indicate *SD*, n = 3; c) Immunofluorescent staining of cultures with varying proportions of SCs for desmin (green) and Hoechst (blue) after 72 h in serum-free myogenic differentiation media; scale bar = 100 μm; d) Fusion indices (nuclei within desmin-stained areas as a proportion of total nuclei) corresponding to c), shown as box plots, n = 4; e) Normalised gene expression in SCs and FAPs at passage 2 (Timepoint 3). Points represent genes, which are coloured if significantly (FDR < 0.05, log-FC > 1) upregulated in FAPs (blue) or SCs (orange). Selected genes are annotated. f) Violin plots showing expression of differentially expressed receptors in SCs and FAPs at passage 2; g) Flow cytometry plots of muscle-derived cells after 72 h in SFGM stained for ITGA5/ITGA7 (left) or CD29/NCAM1 (right) prior to FACS; cells are shown after removal of doublets, CD45+ and JAM-1+ cells; dashed lines indicated gating strategy for sorting; h) Proportion of SCs purified with FACS protocols shown in g) during long-term culture, as measured by flow cytometry; error bars indicate *SD*, n = 3; i) Proportion of SCs during long-term culture on different coatings as indicated, n = 3; j) As i), but for different media formulations as indicated, n = 3.

We therefore sought to adapt the long-term culture conditions, with the aim of preventing FAPs from overgrowing SCs. To inform our approach, we filtered the scRNA-seq data for differentially expressed genes between SCs and FAPs encoding receptors that interact with proteins commonly used as surface coatings or growth factors, revealing a number of interesting candidates (Fig. 4e, f). Sorting SCs using an ITGA7+/ITGA5− strategy significantly decreased FAP contamination as compared to a CD29+/NCAM1+ sorting strategy (used in previous studies^22^), reducing FAP overgrowth after multiple passages (Figs. 4g, h). Culturing SCs contaminated with FAPs on different coatings, we observed that within 3 passages, SC purity was reduced on collagen I and fibronectin (FN; interacting with ITGA5) but remained high on laminin-521 (LN-521; interacting with ITGA7; Fig. 4i). We further adapted SFGM by adding triiodothyronine (T3, 30 nM, ligand for THRA), increasing the concentration of HGF (ligand for c-Met) and removing PDGF-BB (ligand for PDGFRα; Fig. 4j). Each individual adaptation reduced FAP overgrowth over 3 passages, whilst the combination of all three showed no significant reduction in SC proportion over the course of the experiment, and was thus labelled ‘improved SFGM’ (i-SFGM).

### SCs transition between three dynamic states

To further investigate SC heterogeneity in vitro, we filtered out FAPs and reanalysed the remaining single-cell transcriptomes at passage 2 (Timepoint 3). Reclustering identified three subpopulations within SCs, with distinct gene expression profiles suggestive of quiescent, activated, and committed states (similar to those previously observed in murine SCs^14^; Figs. 5a, b). Genes encoding proteins related to focal adhesion and ECM organisation were upregulated in the quiescent state, cell cycle and replication in the activated state, and myogenic differentiation the committed state (Fig. 5c, Supplementary Fig. 5a). These cell states were also observed at passages 5 and 8, but in different ratios (Supplementary Figs. 5b, c), suggesting the possibility of transitioning between states.

**Figure 5:**
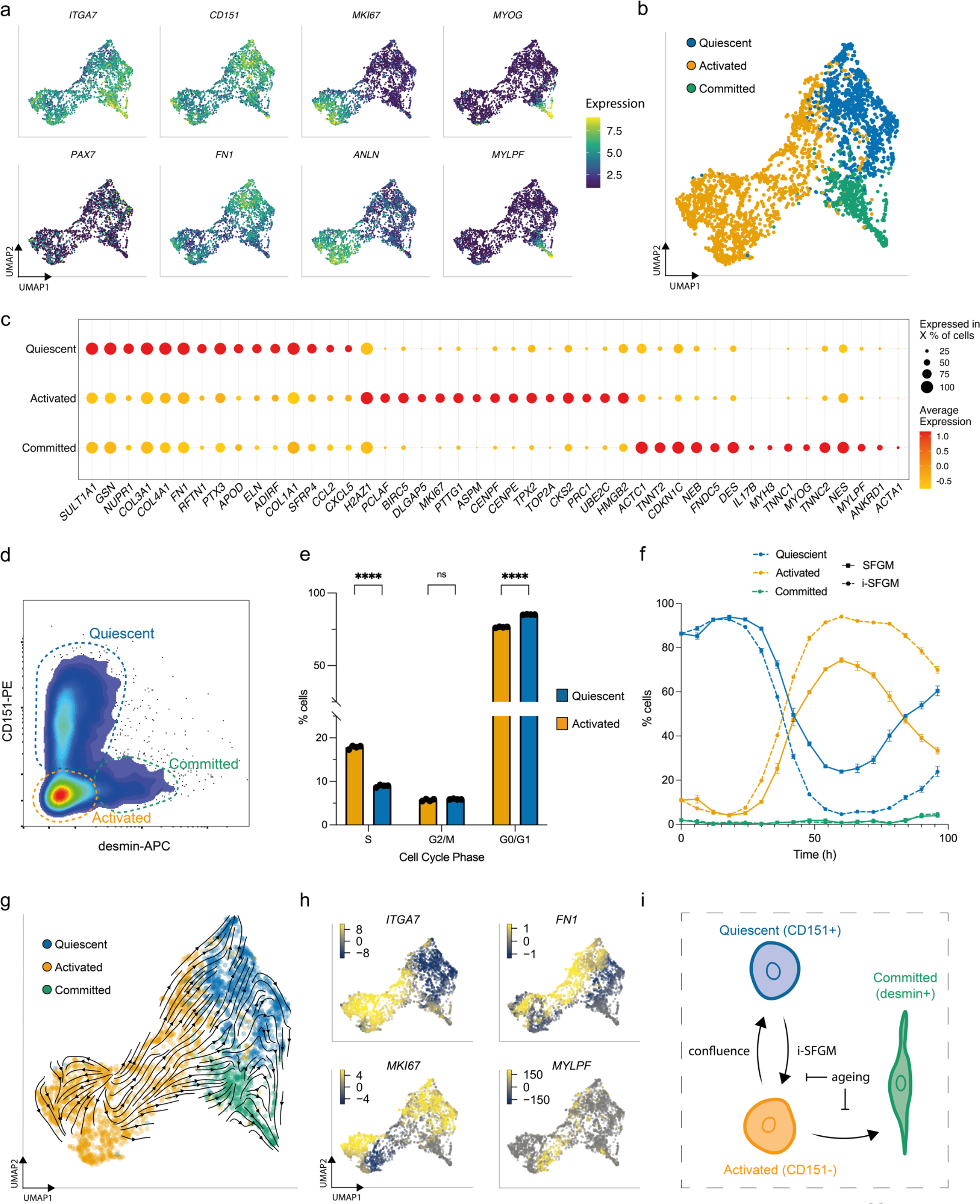
SCs switch dynamically between three cellular states in vitro. a) UMAPs of SCs at passage 2 (Timepoint 3), coloured by expression of indicated genes; b) UMAP as a), but labelled according to SC state; c) Normalised expression of the 15 most upregulated genes, averaged within SCs in each state; d) Flow cytometry plot of purified SCs stained for CD151 and desmin; dashed gates indicate the respective cell states; e) Cell cycle analysis of CD151+ (blue) and CD151- cells (orange) determined via flow cytometry using the gating strategy in Supplementary Fig. 5e; f) Proportion of activated, quiescent and committed SCs over one passage in SFGM or i-SFGM media, as determined via flow cytometry; g) UMAP as b), with averaged RNA velocity vectors embedded; h) UMAPs as a), coloured by RNA velocities of denoted genes; i) Dynamic model of SC states in vitro; arrows indicate direction(s) of proposed transitions between states.

We next established protocols to measure these states in vitro, through analysis of CD151 (upregulated in quiescent SCs^14^) and desmin (upregulated in committed SCs) via flow cytometry (Fig. 5d, Supplementary Fig. 5d). Combining this panel with EdU-staining, we confirmed increased cell cycle activity in activated as compared to quiescent SCs (Fig. 5e; Supplementary Fig. 5e). Investigating the dynamics of these subpopulations during a single passage, we found dynamic conversion between quiescent (CD151+) and activated (CD151−) states, with an initial increase in quiescent SCs followed by a rapid decline and a renewed increase as cells approach confluence (Fig. 5f). The proportion of cells in the committed state remained stable throughout the first 72 h, increasing after 96 h. Investigating over multiple passages revealed that this interchangeability decreased over time (Supplementary Fig. 5f). We observed that i-SFGM increased the proportion of activated SCs compared to SFGM, both within a single passage (Fig. 5f) and over the course of multiple passages (Supplementary Fig. 5f). In addition to transitions between cell states changing over the course of cell aging, transcriptional changes related to increased ECM remodelling and decreased translation were also observed *within* states over time (Supplementary Fig. 6).

Finally, we performed RNA velocity analysis of SCs at passage 2, in order to investigate these cell state transitions in silico. RNA velocity averaged across all genes indicated transitions from the activated to the quiescent state or committed states (Fig. 5g), whilst an increase in expression of cell cycle genes within a subset of quiescent cells suggested they are able to return to the activated state (Fig. 5h). Taken together with in vitro data, this suggests a dynamic, reversible transition between activated and quiescent states, whilst activated cells can also differentiate towards committed myocytes as SCs become confluent (Fig. 5i).

## Discussion

The complexity of muscle tissue arises from the interplay of multiple cell types^8,9^. Here, we have presented an annotated scRNA-seq dataset comprising over 36,000 muscle cellular transcriptomes, from 10 donor cattle, across five timepoints during a cultured beef production process. This dataset gives significant insight into the cellular heterogeneity present in bovine skeletal muscle for the first time, including comprehensive analysis of 11 defined cell types (Fig. 1). Our analysis showed notable similarity between bovine muscle and analogous approaches employed for more well-studied species, with expression of canonical genes for distinct cell types largely conserved between cattle, human and mouse. Comparison of bovine muscle with the corresponding cell cultures shed light on transcriptional changes that occur during the transition to in vitro culture. Notably, genes related to protein synthesis were strongly upregulated (suggesting cell activation), whilst those relating to ECM were downregulated, highlighting the pivotal role of cell-matrix interactions in the muscle niche. However, despite the absence of signalling from the microenvironment, the expression of most canonical marker genes was conserved in vitro (Fig. 2), suggesting that muscle-derived cell cultures are able to recapitulate in vivo behaviour to a large extent.

Differential expression analysis allowed us to identify surface marker panels for physical separation of multiple cell types, including SCs, FAPs, ECs and SMMCs (together representing over 80% of mononucleated cells). Notably, inclusion of ITGA5 in the panel offered significant purity improvements over previous protocols, as this integrin serves as a negative selection marker for SCs, whilst ITGB1/CD29 (previously used as a SC marker^22,23^) is highly expressed by both SCs and FAPs. Whilst we were able to proliferate each of these cell types for multiple passages (Fig. 3), further work is needed to assess the potential of ECs and SMMCs, with respect to their long-term proliferation in the absence of FBS and their capacity to promote myogenesis or adipogenesis, for example in co-culture systems^12,24^. Other low-abundance cell types, such as macrophages, could also offer benefits in terms of tissue remodelling and angiogenesis for structured cultured meat products^25,26^.

Cultured meat production requires an extensive proliferation phase, with many PDs, to achieve required cell yields^5^. This prompts several issues, including reduced growth rates, loss of differentiation capacity, and the possibility of overgrowth of undesired cell types. Transcriptomic analysis indeed revealed that sorted SCs were rapidly overgrown by a small proportion of contaminating FAPs, indicating variable growth dynamics in our initial culture conditions that favour FAP proliferation (Fig. 4). Whilst high levels of animal-derived sera have traditionally been used to control fibroblastic overgrowth in SC cultures^27^, this phenomenon is poorly understood, and in any case is unsuited for cultured meat applications. We used differential expression analysis to inform several improvements to the SC culture conditions, including the switch to an ECM mimic that corresponds to the integrin expression profile of SCs (laminin is known to interact with ITGA7/ITGB1 dimers^28^), and the addition of growth factors, such as HGF, whose receptor expression is specific to SCs^29,30^. Our dataset will also provide insights into design of selective media for other cell types, such as ECs, which is likely complicated by their slow proliferation. Detailed metabolic profiling, which was beyond the scope of this study, could help to further inform the design of feeding strategies that afford efficient and selective growth of desired cell types^31^. It should be noted that our experiments are conducted at lab scale, and that proliferation in bioreactor systems used for upscaled cultured meat production will introduce additional stresses, that could act differentially across cell types^32^. Further cost reduction will also be required, and our dataset can assist in the replacement of growth and attachment factors with peptide alternatives with retained (or superior) cell type selectivities.

Our transcriptomic analysis also revealed remarkable heterogeneity *within* cell types, notably between distinct subpopulations of SCs, that points towards a dynamic equilibrium of quiescent, active and committed states (Fig. 5). Different SC states have previously been proposed, both from dissociated muscle samples^15,33–35^ and in vitro cultures^16,36,37^, although without clear consensus. We identified CD151 (previously observed in human skeletal muscle^13^) as a cell-surface marker for the ‘quiescent’ cluster, although the extent to which the quiescent phenotype we observed in vitro reflects physiological SC behaviour during embryonic development or wound healing remains unclear. We did not distinguish such states in our bovine muscle samples, perhaps because the majority of cells are activated during tissue dissociation^33^. RNA velocity analysis suggests a model in which only activated SCs are able to differentiate, which is congruent with a physiological model of wound healing in which activated SCs either differentiate to form muscle fibres, or return to a quiescent state in order to preserve the ability of the tissue to respond to future regenerative stimuli. However, higher resolution velocity analysis of SCs would be informative, given that we only sampled a snapshot within each passage (when cells were approaching confluency). Indeed, the role that confluency plays in the promotion of differentiation, and the extent to which premature differentiation might negatively affect a proliferative culture, requires further study in the context of a cultured meat bioprocess design. Similarly, whilst increased culture length clearly affects the transitions of SCs between states, it is important to note that we also observed substantial transcriptomic differences between equivalent states at different timepoints (Supplementary Fig. 6), indicating that cellular aging cannot be understood solely in terms of subpopulation ratio changes. Indeed, our data supports the hypothesis that a reduced rate of switching between states could be a characteristic of SC aging or senescence^38,39^. Future transcriptomic experiments, including single-nuclei approaches, could illuminate the formation of cultured muscle tissues during the differentiation of SCs. Furthermore, although not specified in this study, it is likely that similar cellular heterogeneity is present in other cell types, including FAPs, where adipogenic and fibroblastic fate decisions could be critical for cultured fat production^40,41^.

In conclusion, scRNA-seq is a powerful approach to study cell heterogeneity in the context of cultured meat production. Our dataset offers a number of important insights for cell purification and proliferation steps, has led to the development of refined medium conditions, and represents a useful resource for further analysis and improvement of cultured meat bioprocesses.

## Material & Methods

### Cell isolation

Cells were isolated from semitendinosus muscles of 10 Belgian Blue cattle (Supplementary Table 1) by collagenase digestion (CLSAFA, Worthington; 1 h; 37 °C), filtration with 100 μm cell strainers, red blood lysis using a Ammonium-Chloride-Potassium (ACK) lysis buffer, and final filtration with 40 μm cell strainers. Cells from each donor animal were plated as separate cultures in serum-free growth medium^42^ (SFGM, Supplementary Table 3) on fibronectin-coated (4 μg cm^−2^ bovine fibronectin; F1141, Sigma-Aldrich) tissue culture vessels for 72 h prior to FACS, or (where denoted) in growth medium (GM) containing 20% foetal bovine serum (heat-inactivated FBS; 10500-064, Gibco).

### Cell culture

Tissue culture vessels were coated with fibronectin for unsorted cells, ECs, and SMMCs, laminin-521 (0.5 μg cm^−2^, LN521-05, Biolamina) for SCs, or collagen I (0.6 μg cm^−2^, bovine skin collagen; C2124, Sigma-Aldrich) for FAPs for 1 h prior to plating. Unsorted cells, SCs and FAPs were cultured in SFGM (except where otherwise noted), and passaged every 3-4 days upon reaching confluency. ECs and SMMCs were cultured in BioAMF-3 (01-196-1A, Sartorius).

To assess long-term proliferation, cells were cultured as described above for each cell type. Cells were harvested upon approaching confluence, counted, reseeded at 5×10^3^ cm^−2^ and purity assessed at each passage via flow cytometry.

### Single cell RNA-sequencing

#### Cell harvesting

Five timepoints throughout long-term in vitro culture were selected for single-cell RNA sequencing (scRNA-seq), for each of which cells from all genotypes (donor animals) were pooled in equal ratios and washed with 1% BSA in PBS prior to injection. Timepoints 1 to 5 corresponded to unsorted cells directly after isolation (‘Muscle’) and after 72 h (‘passage 0’), and to cultured cells (unsorted cells, SCs and FAPs) at passages 2, 5, and 8 respectively (Fig. 1a, Supplementary Table 1).

#### Library preparation and sequencing

25000 cells were injected for each timepoint into Chromium Single Cell Controller, emulsified with bar-coded gel beads and libraries were constructed following the protocol of the Chromium NextGEM Single Cell 3’ Kit V3.1 (10x Genomics). Quality control of the DNA library was performed on Qiaxcel (QIAgen) and quantified by qPCR using the KAPA SYBR Fast qPCR Master Mix (Illumina). Paired- end single cell 3’ gene expression libraries were sequenced on a Novaseq 6000 System (Illumina) using a NovaSeq S1 flow-cell to a depth of at least 3.5 × 10^8^ reads/timepoint.

#### Data processing and demultiplexing

Raw base call files were demultiplexed using the cellranger mkfastq function (CellRanger 6.0.1). Reads were aligned to Bos Taurus genome (build ARS-UCD1.2) with Ensembl annotations (release 101), using CellRanger count function. Default filtering parameters of CellRanger were applied to obtain a gene expression matrix. Genotypes were deconvolved and assigned to individual cells using demuxlet^43^, based on VCF files previously generated by genotyping of each donor animal with a BovineSNP50 v3 DNA Analysis BeadChip (Illumina).

#### Quality control and normalisation

Across the five timepoints, 36129 cells (90.8%) passed quality control (within 3 median absolute deviations of the median for expressed genes, total counts and percentage mitochondrial genes; Supplementary Fig. 1). Counts for each timepoint were normalised using the sctransform function of Seurat with default parameters^44,45^, regressing out percentage of mitochondrial genes, library size, number of genes, and cell cycle effects.

#### Dimensionality reduction and differential gene expression analysis

Cells were clustered using the FindNeighbors() and FindClusters() functions based on the first 50 principal components and with a resolution of 0.1 respectively. Dimensionality was reduced by uniform manifold approximation and projection (UMAP)^46^ using the first 30 principal components as input, 50 neighbouring points and a minimal distance of 0.1).

Differentially expressed (DE) genes were computed for clusters identified in each timepoint by using the FindAllMarkers(), or between denoted conditions using FindMarkers(), with a log2 fold-change (log-FC) threshold of 1, a false discovery rate (FDR) of 0.05, and expressed in at least 50% of cells. For Timepoint 1, cell types were assigned to each cluster based on DE genes and on predicted phenotypes from FindTransferAnchors()^47^ using analogous murine scRNA-seq data (GSE143437^14^). ECs were further characterised by their expression of signatures derived from the Descartes human genes expression atlas^19^ using the AddModuleScore function. For Timepoint 2, surface markers amongst the DE genes were identified by filtering for genes coding for proteins located in the plasma membrane^48^. GO terms for biological processes (2021) were computed using enrichR^49^. For the analysis of SC cultures, Timepoints 3 to 5 were filtered for genotypes 5, 6, and 10 (Fig. 4), whilst for analysis of SC states these were further filtered by removal of *ITGA5*+ FAP clusters (Fig. 5).

#### RNA velocity analysis

RNA velocities were computed for SCs at Timepoint 3 by generating separate count matrices for spliced and unspliced transcripts using velocyto. The resulting loom files were analysed using scVelo in Python 3.9.12. Transcription kinetics for selected genes were computed, as well as RNA velocity vectors (based on the scVelo dynamical model) across all genes^50,51^.

### Flow cytometry

For cell type identification, unsorted cells were stained for ITGA5, ITGA7, JAM-1 and CD45 (Fig. 3, Supplementary Table 4) for 15 min prior to analysis on a MACSQuant10 Flow Analyzer (Miltenyi Biotec). To determine proportions of SCs and FAPs (Fig. 4), cells were stained for ITGA7 and ITGA5. For analysis of SC states (Fig. 5), cells were stained for CD151 prior to fixation (PFA), permeabilisation (10% saponin), and staining for desmin. Samples were subsequently washed and stained with **α**-mouse-PE secondary antibodies prior to analysis.

### Fluorescent-activated cell sorting (FACS)

Cells were purified by FACS 72 h post-isolation on a MACSQuant Tyto Cell Sorter (Miltenyi Biotec) following antibody staining as described above. SCs, FAPs, SMMCs and ECs were gated as described in Fig. 2. Where noted, cells were gated for NCAM1+/CD29+ (Fig. 4g).

### Immunofluorescent staining

After formaldehyde fixation, cells were permeabilised (0.5% Triton X-100), blocked (2% bovine serum albumin (BSA)) and stained with PDGFRα, Pax7, TEK, and CNN1 (for phenotype confirmation; Fig. 3c) or desmin (for fusion index; Fig. 4c) primary antibodies (see Supplementary Table 4). After washing, cells were incubated with AF488-conjugated secondary antibodies and Hoechst 33342 (Thermo Fisher) prior to imaging on an ImageXpress Pico Automated Cell Imaging System (Molecular Devices).

### Myogenic differentiation assay

Mixed cultures containing SCs and FAPs in varying proportions were seeded on 0.5% Matrigel-coated vessels at a density of 5×10^4^ cm^−2^ in SFGM for 24 h, before switching to serum-free differentiation media (SFDM). After 72 h in SFDM, cells were fixed with 4% formaldehyde, stained for desmin and imaged as previously described. To calculate fusion indices, number of nuclei within desmin-stained areas were divided by total nuclei.

### Cell cycle analysis

For flow cytometric cell cycle analysis, SCs were incubated with EdU for 90 min prior to trypsinisation. Following **α**-CD151-APC staining, Click-iT reaction was performed using the Click-iT EdU Alexa Fluor 488 Flow Cytometry Assay Kit (C10420, Thermo Fisher) according to the supplier’s protocol, followed by **α**-desmin staining as previously described. Finally, the cells were gated into quiescent (DES^−^/CD151^+^) and activated (DES^−^/CD151^−^) and within these, into EdU^+^ (S-Phase), DAPI^low^/EdU^−^ (G0/1-Phase) and DAPI^high^/EdU^−^ (G2-Phase), as shown in Supplementary Fig. 5d.

### Statistical analyses

Statistical analyses were performed using Prism 9 (Graphpad). Pearson correlation of fusion indices with percentages of ITGA7+ cells was computed assuming a normal distribution (Fig. 4d). For comparisons of culture conditions (Figs. 4h-j) and the analysis of cell cycle states in SCs (Fig. 5e), two-way ANOVAs were performed including post-hoc Tukey’s multiple comparison test. Sample replicates consisted of cells from the same donor animal, cultured in separate vessels.

## Data availability

scRNA-seq data is deposited in the NCBI Gene Expression Omnibus database (accession code GSE211428).

## Code availability

Code for scRNA-seq analysis (including RNA velocity calculations) are available from the authors on request.

## Acknowledgements

We would like to thank Latifa Karim, Wouter Coppieters, Alice Mayer and Manon Deckers (GIGA institute, ULiège) for support in the acquisition and analysis of scRNA-seq data, Benjamin Bouchet for advice on immunofluorescent stainings, Christoph Börlin for help with RNA velocity analysis and Dhruv Raina for critical feedback on the manuscript.

## Author contributions

TM, RGJD, LS, LM, RH and MG performed experiments and analysis. MJP and JEF supervised the study. TM and JEF wrote the manuscript with input from all authors.

