## Supplementary Information for "Single-cell analysis of bovine muscle-derived cell types for cultured meat production"

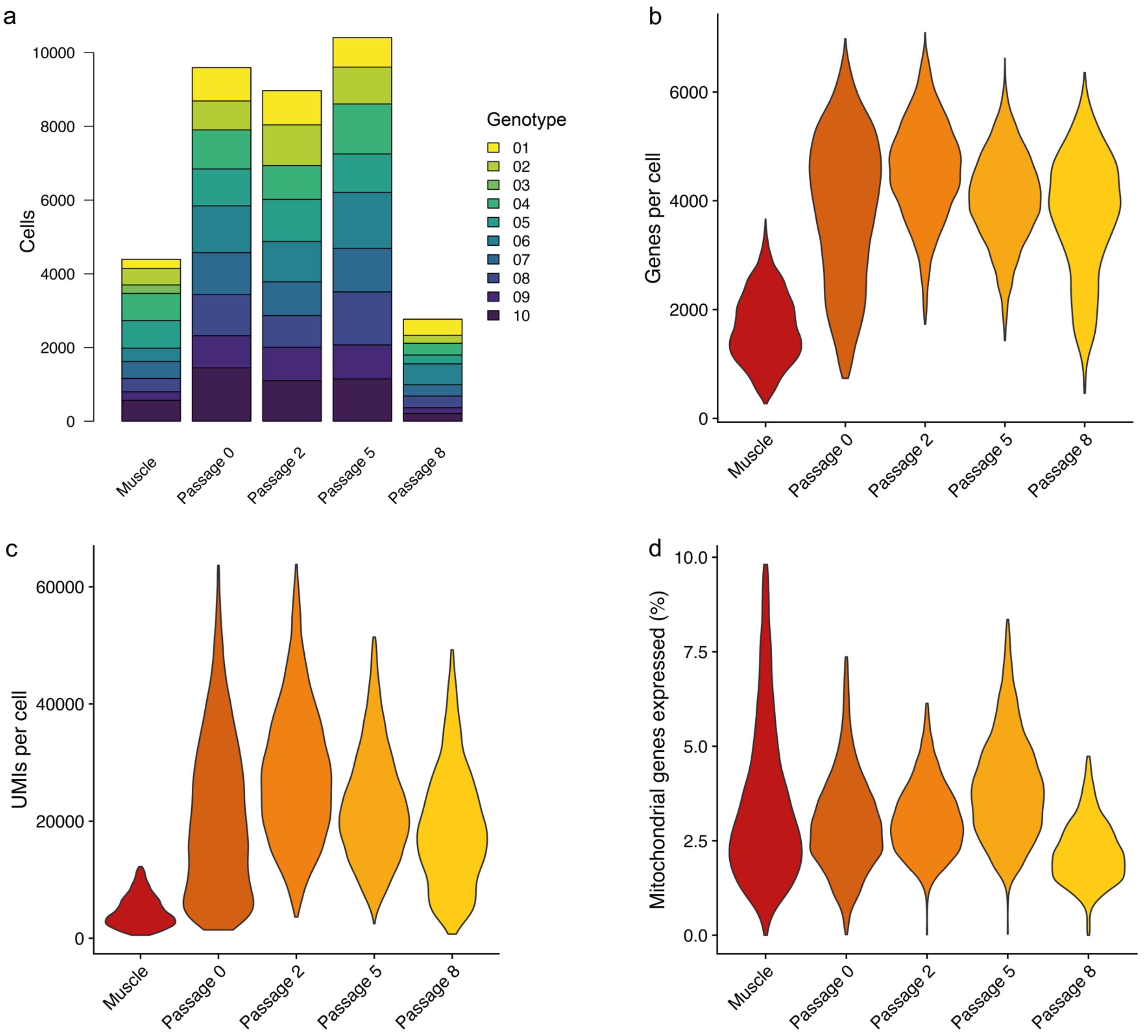


**Supplementary Figure 1: Single-cell RNA-sequencing quality control (related to Figure 1)**

a) Number of cells at each timepoint of the scRNA-seq experiment, coloured by genotype (donor animal);
b) Total number of genes per cell at each timepoint;
c) Total number of unique molecular identifiers (UMIs) per cell at each timepoint;
d) Percentage of reads assigned to mitochondrial genes at each timepoint.

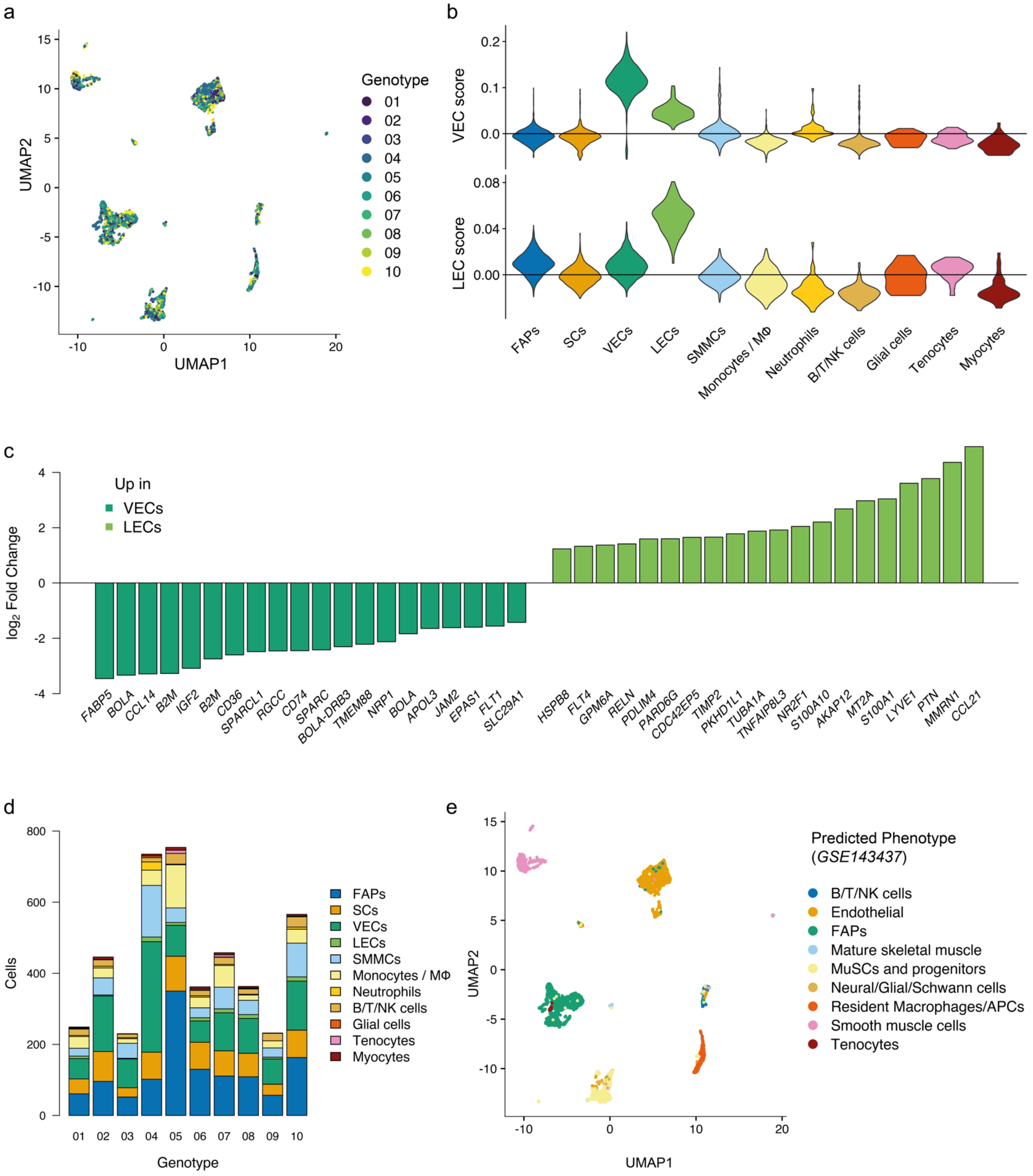


**Supplementary Figure 2: Identification of cell types in the bovine muscle niche (related to Figure 1).**

a) UMAP of bovine muscle cells coloured by genotype;
b) Averaged expression of *Descartes'* gene signature for vascular (top) and lymphatic endothelial (bottom) marker genes per cluster;
c) Fold changes of the 20 most differentially expressed genes between vascular endothelial cells (VECs, dark green) and lymphatic endothelial cells (LECs, light green);
d) Number of cells per genotype at Timepoint 1, coloured by respective cell type;
e) UMAP of bovine muscle cells coloured by predicted phenotype, based on gene expression profiles of murine muscle cells (from *GSE143437*).

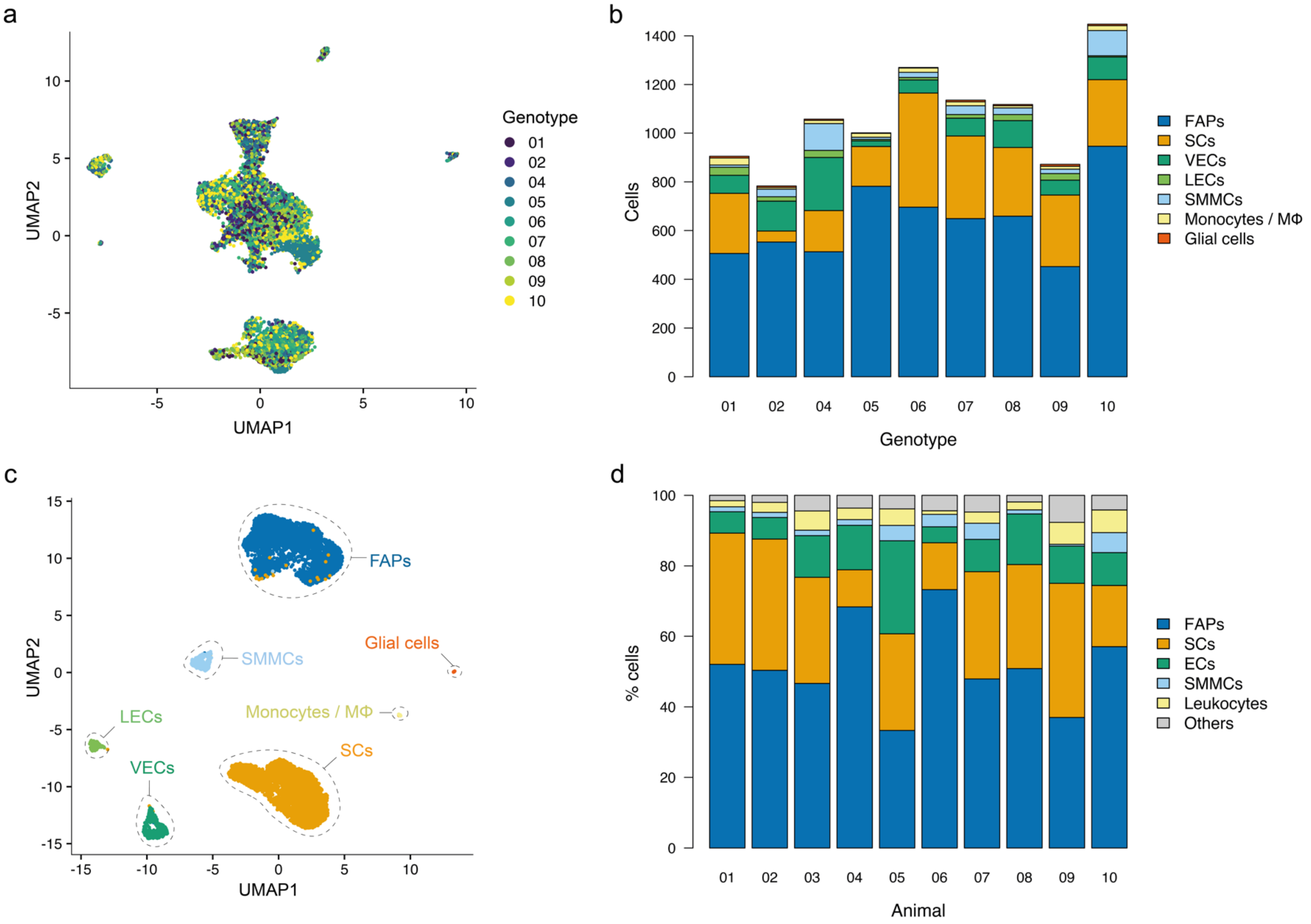


**Supplementary Figure 3: Adherent cell types from bovine muscle niche after 72 h in vitro (related to Figure 2).**

a) UMAP of adherent cell fraction after 72 h in SFGM, coloured by genotype;
b) Number of cells per genotype at Timepoint 2, coloured by respective cell type;
c) UMAP of adherent cell fraction after 72 h in serum-containing GM, coloured by respective phenotype;
d) Percentage of cell types in 10 donor animals, measured via flow cytometry based on gating strategy shown in Fig. 2f.

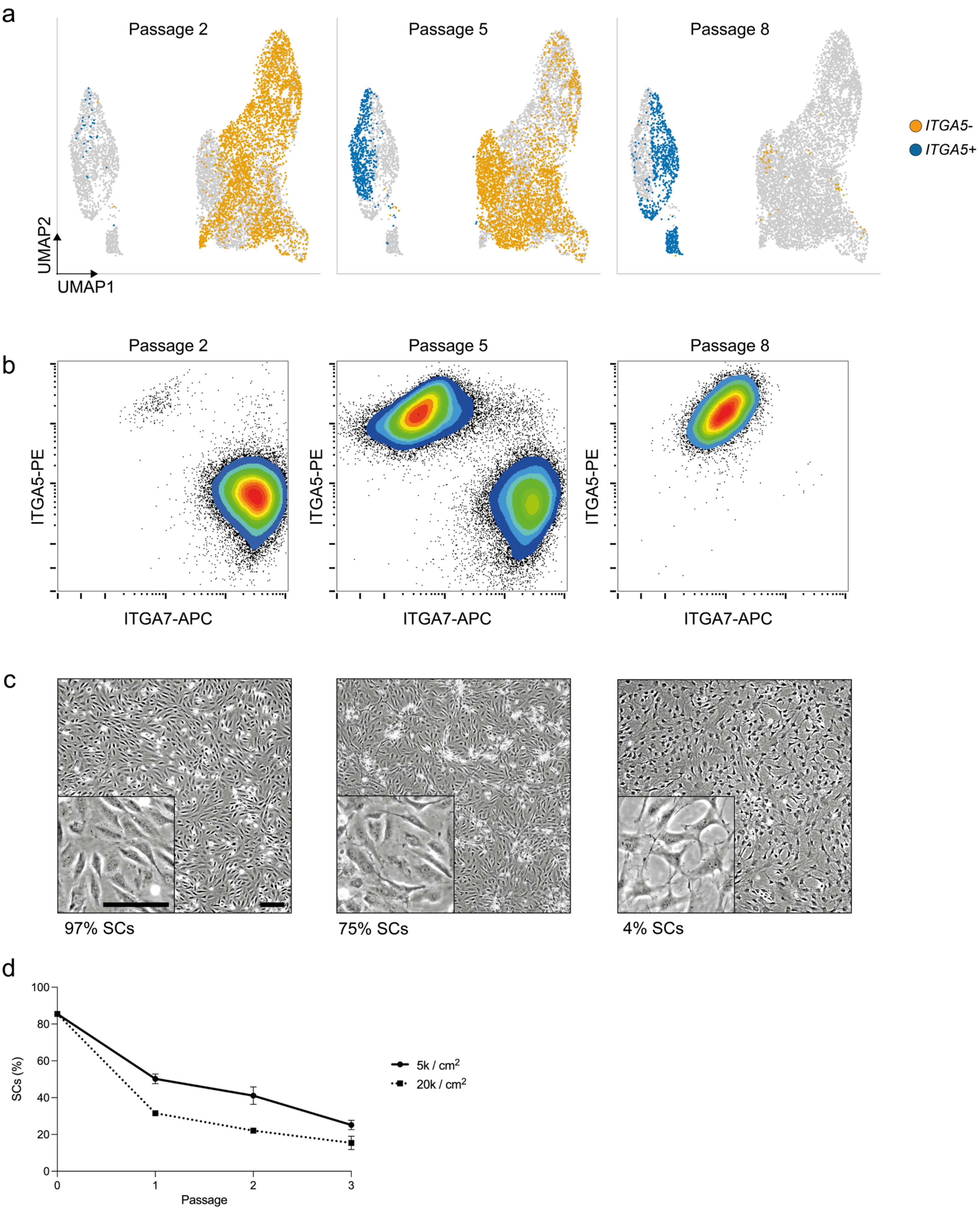
a
Passage 2
Passage 5
Passage 8
ITGA5-
ITGA5+
UMAP2
UMAP1
b
Passage 2
Passage 5
Passage 8
ITGA5-PE
ITGA7-APC
c
97% SCs
75% SCs
4% SCs
d
SCs (%)
100
80
60
40
20
0
0
1
2
3
Passage
5k / cm2
20k / cm2

**Supplementary Figure 4: Overgrowth of SCs by FAPs during long-term cultivation (related to Figure 4).**

a) UMAPs of sorted SCs at passages 2 (Timepoint 3, left), 5 (Timepoint 4, centre) and 8 (Timepoint 5, right), clusters coloured for expression of *ITGA5*;
b) Flow cytometry plots of sorted SCs at passages 2 (left), 5 (centre) and 8 (right);
c) Brightfield images of heterogeneous cultures with denoted ITGA7+ percentages; scale bar = 100 µm;
d) Proportion of SCs resulting from passaging at different seeding densities, as measured via flow cytometry; error bars indicate *SD*, n = 4.

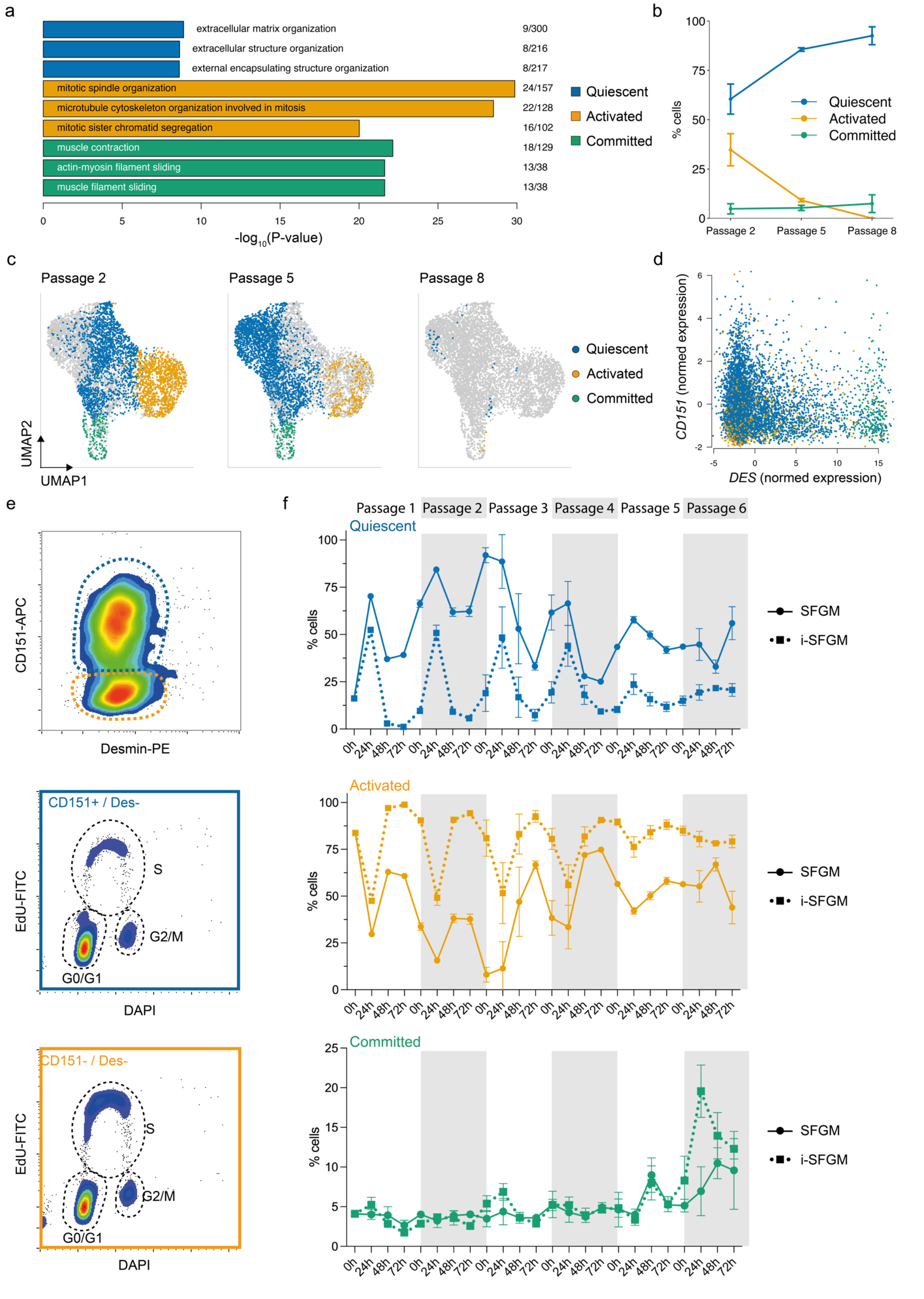

a
extracellular matrix organization
9/300
extracellular structure organization
8/216
external encapsulating structure organization
8/217
mitotic spindle organization
24/157
microtubule cytoskeleton organization involved in mitosis
22/128
mitotic sister chromatid segregation
16/102
muscle contraction
18/129
actin-myosin filament sliding
13/38
muscle filament sliding
13/38
Quiescent
Activated
Committed
-log10(P-value)
b
% cells
Quiescent
Activated
Committed
Passage 2
Passage 5
Passage 8
c
Passage 2
Passage 5
Passage 8
Quiescent
Activated
Committed
UMAP2
UMAP1
d
CD151 (normed expression)
DES (normed expression)
e
CD151-APC
Desmin-PE
CD151+ / Des-
EdU-FITC
S
G2/M
G0/G1
DAPI
CD151- / Des-
EdU-FITC
S
G2/M
G0/G1
DAPI
f
Passage 1
Passage 2
Passage 3
Passage 4
Passage 5
Passage 6
Quiescent
Activated
Committed
% cells
SFGM
i-SFGM
0h
24h
48h
72h

**Supplementary Figure 5: Three dynamic states identified within purified SCs (related to Figure 5).**

a) Top three most significantly enriched GO terms corresponding to upregulated genes genes in quiescent, activated, and committed SCs;
b) Proportions of quiescent, activated and committed SCs at passages 2, 5 and 8, as determined via scRNA-seq;
c) Combined UMAPs showing SCs at passages 2 (left), 5 (centre) and 8 (right), coloured according to cell state;
d) Expression of *CD151* and *DES* in SCs at passage 2; cells are coloured by state;
e) Gating strategy for cell cycle analysis via flow cytometry; SCs were stained for CD151 and desmin (top); CD151+ (blue, centre) and CD151- (orange, bottom) cells were assigned to cell cycle phases as indicated by dotted gates;
f) Proportion of quiescent (top), activated (centre) and committed (right) SCs in SFGM and improved SFGM media over the course of three passages, determined every 24 h via flow cytometry.

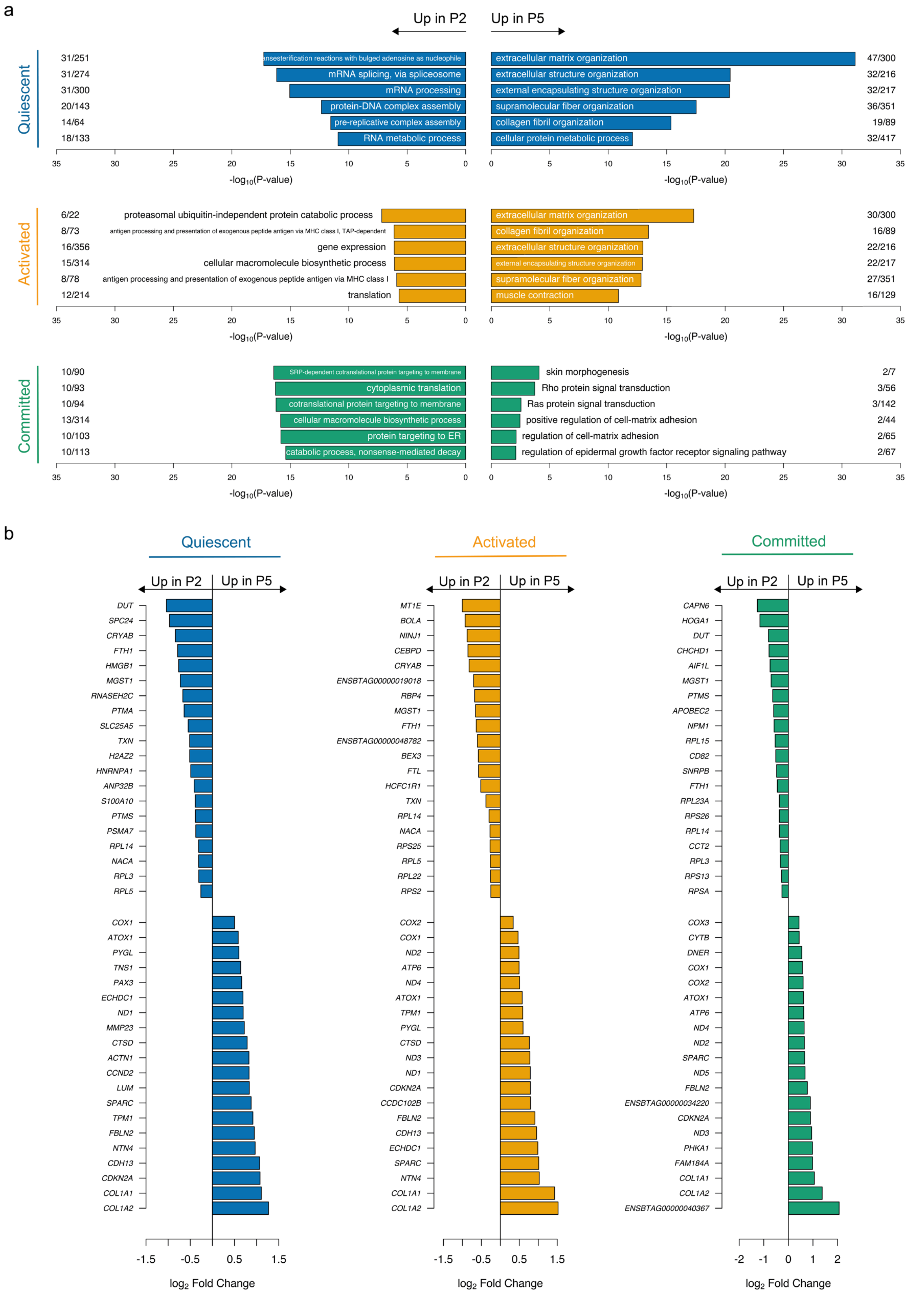

a
Up in P2
Up in P5
Quiescent
31/251 ansesterification reactions with bulged adenosine as nucleophile
31/274 mRNA splicing, via spliceosome
31/300 mRNA processing
20/143 protein-DNA complex assembly
14/64 pre-replicative complex assembly
18/133 RNA metabolic process
extracellular matrix organization 47/300
extracellular structure organization 32/216
external encapsulating structure organization 32/217
supramolecular fiber organization 36/351
collagen fibril organization 19/89
cellular protein metabolic process 32/417
-log10(P-value)
Activated
6/22 proteasomal ubiquitin-independent protein catabolic process
8/73 antigen processing and presentation of exogenous peptide antigen via MHC class I, TAP-dependent
16/356 gene expression
15/314 cellular macromolecule biosynthetic process
8/78 antigen processing and presentation of exogenous peptide antigen via MHC class I
12/214 translation
extracellular matrix organization 30/300
collagen fibril organization 16/89
extracellular structure organization 22/216
external encapsulating structure organization 22/217
supramolecular fiber organization 27/351
muscle contraction 16/129
Committed
10/90 SRP-dependent cotranslational protein targeting to membrane
10/93 cytoplasmic translation
10/94 cotranslational protein targeting to membrane
13/314 cellular macromolecule biosynthetic process
10/103 protein targeting to ER
10/113 catabolic process, nonsense-mediated decay
skin morphogenesis 2/7
Rho protein signal transduction 3/56
Ras protein signal transduction 3/142
positive regulation of cell-matrix adhesion 2/44
regulation of cell-matrix adhesion 2/65
regulation of epidermal growth factor receptor signaling pathway 2/67
b
Quiescent
Activated
Committed
Up in P2
Up in P5
DUT, SPC24, CRYAB, FTH1, HMGB1, MGST1, RNASEH2C, PTMA, SLC25A5, TXN, H2AZ2, HNRNPA1, ANP32B, S100A10, PTMS, PSMA7, RPL14, NACA, RPL3, RPL5, COX1, ATOX1, PYGL, TNS1, PAX3, ECHDC1, ND1, MMP23, CTSD, ACTN1, CCND2, LUM, SPARC, TPM1, FBLN2, NTN4, CDH13, CDKN2A, COL1A1, COL1A2
MT1E, BOLA, NINJ1, CEBPD, CRYAB, ENSBTAG00000019018, RBP4, MGST1, FTH1, ENSBTAG00000048782, BEX3, FTL, HCFC1R1, TXN, RPL14, NACA, RPS25, RPL5, RPL22, RPS2, COX2, COX1, ND2, ATP6, ND4, ATOX1, TPM1, PYGL, CTSD, ND3, ND1, CDKN2A, CCDC102B, FBLN2, CDH13, ECHDC1, SPARC, NTN4, COL1A1, COL1A2
CAPN6, HOGA1, DUT, CHCHD1, AIF1L, MGST1, PTMS, APOBEC2, NPM1, RPL15, CD82, SNRPB, FTH1, RPL23A, RPS26, RPL14, CCT2, RPL3, RPS13, RPSA, COX3, CYTB, DNER, COX1, COX2, ATOX1, ATP6, ND4, ND2, SPARC, ND5, FBLN2, ENSBTAG00000034220, CDKN2A, ND3, PHKA1, FAM184A, COL1A1, COL1A2, ENSBTAG00000040367
log2 Fold Change

**Supplementary Figure 6: Aging effects within SC subpopulations (related to Figure 5).**

a) Most significantly enriched GO-terms corresponding to the 200 most upregulated genes between passage 2 (left) and passage 5 (right) in quiescent (blue), activated (orange), and committed (green) SCs;
b) Fold changes of the 20 most up- and downregulated genes in quiescent (blue), activated (orange), and committed (green) SCs.

**Supplementary Table 1: Experimental overview.**

| Animal | Breed | Sex | Age | Sorted population | Number of cells | | | | |
|---|---|---|---|---|---|---|---|---|---|
| | | | | | Timepoint 1 | Timepoint 2 | Timepoint 3 | Timepoint 4 | Timepoint 5 |
| | | | | | Muscle | P0 | P2 | P5 | P8 |
| 01 | Belgian Blue | Female | Adult | Unsorted | 249 | 905 | 927 | 801 | 443 |
| 02 | Belgian Blue | Female | Calf | Unsorted | 446 | 783 | 1102 | 998 | 216 |
| 03 | Belgian Blue | Female | Adult | N/A | 230 | - | - | - | - |
| 04 | Belgian Blue | Female | Calf | FAPs | 735 | 1058 | 917 | 1355 | 317 |
| 05 | Belgian Blue | Female | Calf | SCs | 754 | 1001 | 1144 | 1041 | 239 |
| 06 | Belgian Blue | Male | Adult | SCs | 362 | 1270 | 1092 | 1522 | 567 |
| 07 | Belgian Blue | Female | Calf | FAPs | 458 | 1136 | 915 | 1177 | 307 |
| 08 | Belgian Blue | Male | Adult | FAPs | 363 | 1118 | 863 | 1442 | 316 |
| 09 | Belgian Blue | Female | Adult | Unsorted | 232 | 872 | 905 | 922 | 159 |
| 10 | Belgian Blue | Male | Calf | SCs | 566 | 1448 | 1101 | 1147 | 208 |
| | | | | Total: | **4395** | **9591** | **8966** | **10405** | **2772** |
| | | | | | | | | Overall total: | **36129** |

**Supplementary Table 2: Upregulated genes of identified cell types in bovine muscle (attached as tsv-file).**

**Supplementary Table 3: Media formulations.**

| # | Component | Reference | Concentration |
|---|---|---|---|
| **Serum-free growth medium (SFGM)** | | | |
| 1 | DMEM/F-12 | 21331-020, Gibco | |
| 2 | α-linolenic acid | L2376, Sigma Aldrich | 1.0 μg $ml^{-1}$ |
| 3 | FGF-2 | 100-18B, Peprotech | 10 ng $ml^{-1}$ |
| 4 | Human Serum Albumin | Rc HA NW20, Richcore Lifesciences | 5.0 mg $ml^{-1}$ |
| 5 | HGF | 100-39H, Peprotech | 5 ng $ml^{-1}$ |
| 6 | Hydrocortisone | H0135, Sigma Aldrich | 36 μg $ml^{-1}$ |
| 7 | IGF-1 | 100-11, Peprotech | 100 ng $ml^{-1}$ |
| 8 | IL-6 | 200-06, Peprotech | 20 ng $ml^{-1}$ |
| 9 | ITSE | 00-101, biogems | 1% |
| 10 | GlutaMax | 35050-061, Gibco | 1% |
| 11 | Glucose | G7021, Sigma Aldrich | 17.7 mM |
| 12 | L-ascorbic acid 2-phosphate (Vitamin C) | A8960, Sigma Aldrich | 155 μM |
| 13 | PDGF-BB | 100-14B, Peprotech | 10 ng $ml^{-1}$ |
| 14 | Penicillin/Streptomycin/Amphotericin (PSA) | 17-745E, Lonza | 1% |
| 15 | VEGF | 100-20 Peprotech | 10 ng $ml^{-1}$ |
| **Growth medium (GM)** | | | |
| 1 | Ham's F-10 Nutrient Mix | 31550-023, Gibco | |
| 2 | Fetal Bovine Serum, heat inactivated (FBS) | 10082147, Gibco | 20% |
| 3 | FGF-2 | 100-18B, Peprotech | 5 ng $ml^{-1}$ |
| 4 | PSA | 17-745E, Lonza | 1% |
| **Serum-free differentiation medium (SFDM)** | | | |
| 1 | DMEM/F-12 | 21311-020, Gibco | |
| 2 | EGF-1 | AF-100-15, Peprotech | 10 ng $ml^{-1}$ |
| 3 | Human Serum Albumin | Rc HA NW20, Richcore Lifesciences | 0.5 mg $ml^{-1}$ |
| 4 | ITSE | 00-101, biogems | 2% |
| 5 | L-ascorbic acid 2-phosphate (Vitamin C) | A8960, Sigma Aldrich | 40 μM |
| 6 | Lysophosphatidic acid (LPA) | L7260, Sigma Aldrich | 1 μM |
| 7 | MEM Amino Acids Solution | 11130-051, ThermoFisher | 0.50% |
| 8 | $NaHCO_3$ | P2256, Sigma Aldrich | 6.5 mM |
| 9 | PSA | 17-745E, Lonza | 1% |
| 10 | Soy hydrolysates | 58903C, Merck | 1% |

**Supplementary Table 4: Antibodies used in this study.**

| Target | Colour | Source | Dilution | Reference | Application |
|---|---|---|---|---|---|
| Calponin-1 | - | Abcam | 1:400 | ab46794 | IF |
| CD151 | APC | Miltenyi Biotec | 1:50 | 130-103-664 | Flow |
| CD29 | APC | BioLegend | 1:20 | B247653 | Flow |
| CD45-Ro | APC-Vio770 | Miltenyi Biotec | 1:50 | 130-114-083 | Flow |
| Desmin | - | Sigma Aldrich | 1:1000 | D1033 | Flow; IF |
| ITGA5 | PE | Miltenyi Biotec | 1:50 | 130-110-532 | Flow |
| ITGA7 | APC | Miltenyi Biotec | 1:50 | 130-123-833 | Flow |
| JAM-1 / F11R | PE-Vio770 | Miltenyi Biotec | 1:50 | 130-109-484 | Flow |
| NCAM1 | PE | BD Biosciences | 1:20 | 335826 | Flow |
| Pax7 | - | Developmental Studies Hybridoma Bank | 1:100 | Cat# PAX7 | IF |
| PDGFRα | - | Abcam | 1:200 | ab203491 | IF |
| TEK / Tie2 | - | BioLegend | 1:100 | 334202 | IF |
| mouse | AF488 | Invitrogen | 1:250 | A-11029 | IF |
| mouse | PE | Miltenyi Biotec | 1:250 | 30-119-684 | IF |
| rabbit | AF488 | Invitrogen | 1:250 | A-11034 | IF |
